# Relation Extraction for Diet, Non-Communicable Disease and Biomarker Associations (RECoDe): A CoDiet study

**DOI:** 10.64898/2026.03.03.709244

**Authors:** Donghee Choi, Yajie Gu, Kevin Zong, Antoine D. Lain, Dimitrios Zaikis, Thomas Rowlands, Marek Rei, The CoDiet consortium, Tim Beck, Joram M. Posma

## Abstract

Diet plays a critical role in human health, with growing evidence linking dietary habits to disease outcomes. However, extracting structured dietary knowledge from biomedical literature remains challenging due to the lack of dedicated relation extraction datasets. To address this gap, we introduce RECoDe, a novel relation extraction (RE) dataset designed specifically for diet, disease, and related biomedical entities. RECoDe captures a diverse set of relation types, including a broad spectrum of positive association patterns and explicit negative examples, with over 5,000 human-annotated instances validated by up to five independent annotators. Furthermore, we benchmark various natural language processing (NLP) RE models, including BERT-based architectures and enhanced prompting techniques with locally deployed large language models (LLMs) to improve classification performance on underrepresented relation types. The best performing model gpt-oss-20B, a local LLM, achieved an F1-score of 64% for multi-class classification and 92% for binary classification using a hierarchical prompting strategy with a separate reflection step built in. To demonstrate the practical utility of RECoDe, we introduce the Contextual Co-occurrence Summarisation (Co-CoS) framework, which aggregates sentence-level relation extractions into document-level summaries and further integrates evidence across multiple documents. Co-CoS produces effect estimates consistent with established dietary knowledge, demonstrating its validity as a general framework for systematic evidence synthesis.

**Availability:** The code, models, and data will be made freely available upon acceptance.

## Introduction

Diet plays a fundamental role in human health, impacting both the prevention and progression of non-communicable diseases (NCDs). Dietary factors shape disease outcomes through well-defined relationships, including their role in regulating blood pressure (1), modulating lipid metabolism (2), influencing weight management (3), and contributing to metabolic changes in ageing (4).

Meanwhile, systematic reviews play a crucial role in synthesising dietary evidence, providing a rigorous and transparent approach to evaluating unstructured biomedical literature while minimising bias and informing research and policy decisions (5). However, the sheer volume of unstructured data makes traditional systematic reviews slow and labour-intensive. Recent advancements in natural language processing (NLP) offer promising, yet still evolving, approaches to automating systematic review processes, enhancing efficiency in screening and extracting relevant evidence (6).

Among these developments, biomedical information extraction (IE) methods facilitate the identification of key entities and relations, forming the foundation for structuring knowledge from unstructured text (7). Building on such approaches, biomedical knowledge graphs (KGs) provide a scalable framework for integrating diverse insights and uncovering complex associations (8). However, diet-disease relations remain under-explored, necessitating dedicated datasets and tailored methodologies to bridge this gap.

Despite advances in biomedical relation extraction (RE), existing datasets predominantly focus on drug-disease and gene-disease interactions, leaving dietary relationships significantly underrepresented. This limitation prevents NLP models from effectively capturing the nuanced effects of diet on health. The lack of structured datasets tailored for diet-disease RE hinders the development of models capable of systematically extracting and analysing these relationships.

To bridge this gap, we introduce RECoDe, a high-quality, manually annotated dataset specifically designed for diet-disease relation extraction. Building upon previous efforts in diet-related corpus creation (9), we expand this work by developing a dataset that captures fine-grained relationships between diet, disease, and related bio-entities. Understanding these complex associations is essential for assessing disease risk and outcomes, providing a structured foundation for dietary impact analysis (10).

To evaluate RECoDe, we benchmark multiple NLP models, including BERT-based architectures (11), enhanced prompting techniques with locally deployed large language models (LLMs), and compare our method against the recent DiMB-RE framework (12), which is a benchmark dataset and framework for extracting diet–microbiome relations from biomedical literature. Our analysis highlights the advantages and challenges of leveraging LLMs for diet-disease RE, particularly in handling underrepresented relation types.

Our key contributions include:

- The first relation extraction dataset for diet-disease relation extraction, enabling structured dietary knowledge discovery.
- A detailed annotation schema that captures complex dietary interactions with high granularity.
- Extensive benchmarking using transformer-based models and LLM-guided approaches to improve classification performance on underrepresented relation types.

By providing a structured foundation for dietary relation extraction, RECoDe paves the way for more accurate and scalable biomedical NLP applications in nutrition science. We anticipate that our dataset will serve as a valuable resource for computational health research, facilitating deeper insights into the role of diet in disease mechanisms.

## Data and Methods

### RECoDe Dataset

Existing biomedical relation extraction datasets primarily focus on specific molecular interactions, such as drug-disease (13), gene-disease (14), and protein-protein interactions (15). One of the most comprehensive biomedical RE datasets, BioRED, covers various entity types, including disease, gene, and chemical (16). However, even BioRED lacks dietary components, limiting its applicability to nutrition research. We use the CoDiet corpus dataset (9) and from this resource, we selected 109 full-text documents from PubMed Central (PMC), comprising 30 for training, 30 for validation, and 49 for testing. For our experiments, we rely exclusively on the gold-standard named entity recognition (NER) annotations provided in the original corpus. The details of the annotation process and guidelines can be found in the CoDiet corpus paper (9).

We conducted two relation annotation phases: the main annotation period and the adjudication period (see Figure 1). During the first annotation period, one annotator was instructed to select as many candidate relation pairs as possible across the entire document. In the second period, two main annotators independently annotated the same set of 5,014 relation instances. The inter-annotator agreement (IAA) score between the two annotators was 0.919.

**Fig. 1.**
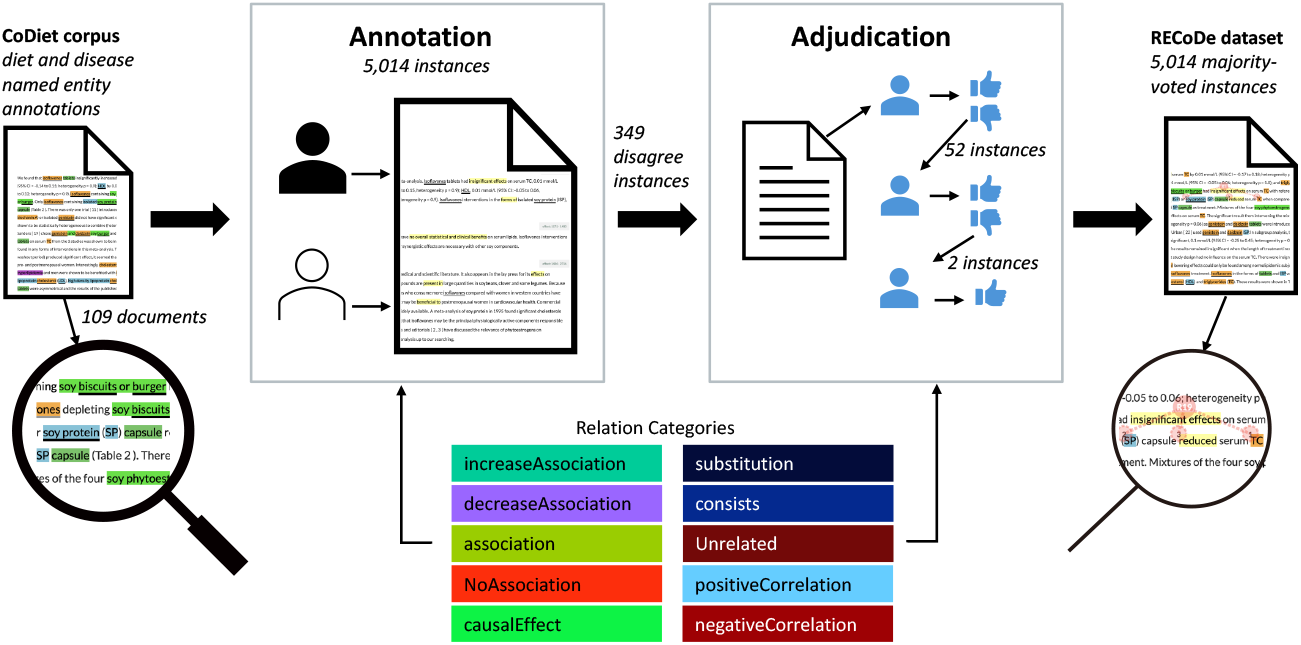
Overview of the annotation and adjudication process in the CoDiet corpus. A total of 5,014 majority-voted instances were included, with 349 disagreements adjudicated across 109 documents.

In the subsequent adjudication period, we ensured that all instances were finalised through majority voting. Since the task involves 10 relation classes, annotators occasionally selected different labels for the same instance. For such cases, additional annotators were involved to resolve the disagreements. Specifically, out of 5,014 instances, 349 cases required adjudication. The first additional annotator resolved 349 disagreements, leaving 52 unresolved. A second annotator addressed these 52, leaving 2 remaining cases, which were finally adjudicated by a third annotator. So in total, more than 3 people agreed to the final dataset. The test set will be hosted on Codabench upon publication to allow benchmarking.

Table 1 compares the distribution of relation types in major biomedical RE datasets. Traditional corpora (e.g., AIMed, ADE, DDI, N2C2, RENET2) are dominated by a small number of relation categories, often focusing on drug–adverse event or protein–protein interactions, and typically lack explicit negative relations. BioRED extends coverage to correlations, but does not capture dietary knowledge. In contrast, RECoDe covers a diverse set of relations, including *association* (19.8%), *causalEffect* (11.1%), *increase/decrease association* (25.8%), and explicit negatives such as *NoAssociation* (6.6%) and *Unrelated* (14.4%). Detailed statistics of entity pair distributions across the train, validation, and test splits are provided in Supplementary Figure S1. These results confirm that RECoDe covers a broad range of diet–biomedical entity combinations.

**Table 1.** Relation Types in Biomedical Datasets. − indicates its absence of a relation type in the dataset. Negative relations are further categorized into NoAssociation and Unrelated.

| Dataset | incAsso | decAsso | asso | causal | subst | consist | posCorr | negCorr | NoAssoc | Unrela | Total |
| --- | --- | --- | --- | --- | --- | --- | --- | --- | --- | --- | --- |
| AIMed (17) | - | - | 100% | - | - | - | - | - | - | - | 1,101 |
| BioInfer (18) | - | - | 19% | - | - | 21% | 54% | 6% | - | - | 2,662 |
| EMU (19) | - | - | 100% | - | - | - | - | - | - | - | 179 |
| ADE (20) | - | - | 4% | 96% | - | - | - | - | - | - | 7,100 |
| DDI (21) | - | - | 59% | 41% | - | - | - | - | - | - | 5,028 |
| N-array (22) | - | - | 100% | - | - | - | - | - | - | - | 3,462 |
| N2C2 (23) | - | - | 97% | 3% | - | - | - | - | - | - | 59,810 |
| DrugProt (24) | 17% | 29% | 54% | - | - | - | - | - | - | - | 24,526 |
| RENET2 (25) | - | - | 100% | - | - | - | - | - | - | - | 51,642 |
| BioRED (16) | - | - | 55% | - | - | - | 27% | 18% | - | - | 6,503 |
| DIMB-RE (12) | 34% | 20% | 7% | 23% | - | 5% | 7% | 4% | - | - | 4,206 |
| **RECoDe** | **12.1%** | **13.7%** | **19.8%** | **11.1%** | **0.4%** | **16.0%** | **3.4%** | **2.4%** | **6.6%** | **14.4%** | **5,014** |

### Experiments

The primary aim of our experiments is to identify the most effective and practically feasible approaches for RECoDe relation extraction task within realistic academic resource constraints. Accordingly, we evaluate both supervised and generative approaches for the RECoDe relation extraction task. For the **supervised setting**, we employ traditional BERT-like models trained using standard supervised learning on our annotated training dataset. In contrast, the **generative setting** explores recent advances in LLMs that can infer relations through prompting rather than parameter fine-tuning. Detailed descriptions of all evaluated models and evaluation protocols, including architecture, parameter sizes, deployment configurations, and performance metrics, are provided in the Models section of the Supplementary Materials.

### Prompt and Inference Strategy

All prompt tuning experiments were conducted using the training set, which we regard as a form of supervised learning aimed at improving generalisability. Firstly, we applied a prompting approach using a set of well-established techniques, including role-play prompting (26), guideline-based prompting, and example-augmented prompting, which have shown effectiveness in biomedical QA tasks (27). We propose a decision-flow framework, illustrated in Fig. 2. The motivation arises from the observation that most local LLMs struggle with limited context length. To address this, we design a multi-stage decision process inspired by decision trees, with a structured workflow resembling recent agentic workflow reasoning strategies (28). In the decision-making process, the model distinguishes between positive relations and negative cases (*NoAssociation* and *Unrelated*). We found that providing more fine-grained definitions of *NoAssociation* and *Unrelated* improved performance compared to a simple binary classification.

**Fig. 2.**
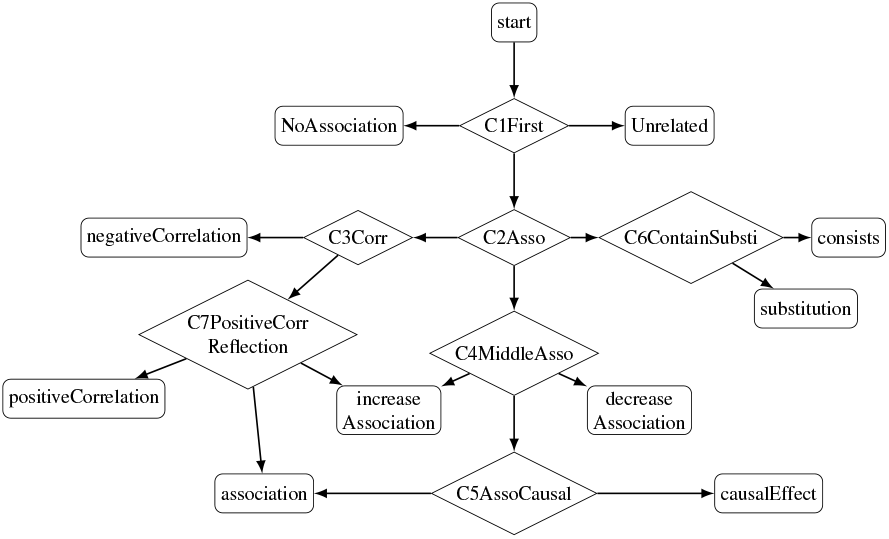
Decision flow of `predict:` staged classifiers (C1–C7) and branching on `majority_pred`.

If the input is classified as a positive relation, the next stage divides it into three categories: *correlation, association*, and *contains*. Correlations are further separated into *positiveCorrelation* and *negativeCorrelation*. Associations are classified into *increaseAssociation* and *decreaseAssociation*. Finally, the *contains* branch is divided into *consists* and *substitution*. All prompts are presented in the Prompts section of the Supplementary Materials. Each prompt begins with a system role-play instruction adapting LLMs to domain-specific tasks. And we provide detailed explanations of the answer classes. We further include guidelines refined through human trials on the training dataset, highlighting notable “dos and don’ts” such as explicitly stating that NoAssociation corresponds to “no effect”. To handle cases that are difficult to describe with textual guidelines alone, we add clear and distinguishable examples. Finally, we incorporate fine-grained instructions such as “Answer (A) Yes ONLY IF…” to enforce precise decision rules.

We also applied a specialised reflection step (29) for the *positiveCorrelation* class, which is designed to allow the LLM to revise its initial prediction particularly for challenging or ambiguous cases. During initial trials with the training set, we observed that LLMs frequently confused *positiveCorrelation* with *increaseAssociation* or *association*.

### Contextual Co-occurrence Summarisation

We define the concept of <u>Co</u>ntextual <u>C</u>o-<u>o</u>ccurrence <u>S</u>ummarisation (CoCoS) as a means to combine the evidence found across different documents for the same entity-pair. Multiple extracted relations for the same entity pair within a document are aggregated into a single document-level relation using the rule-based procedure described in Algorithm 1.

#### Algorithm 1

CoCoS: Within-document relation aggregation

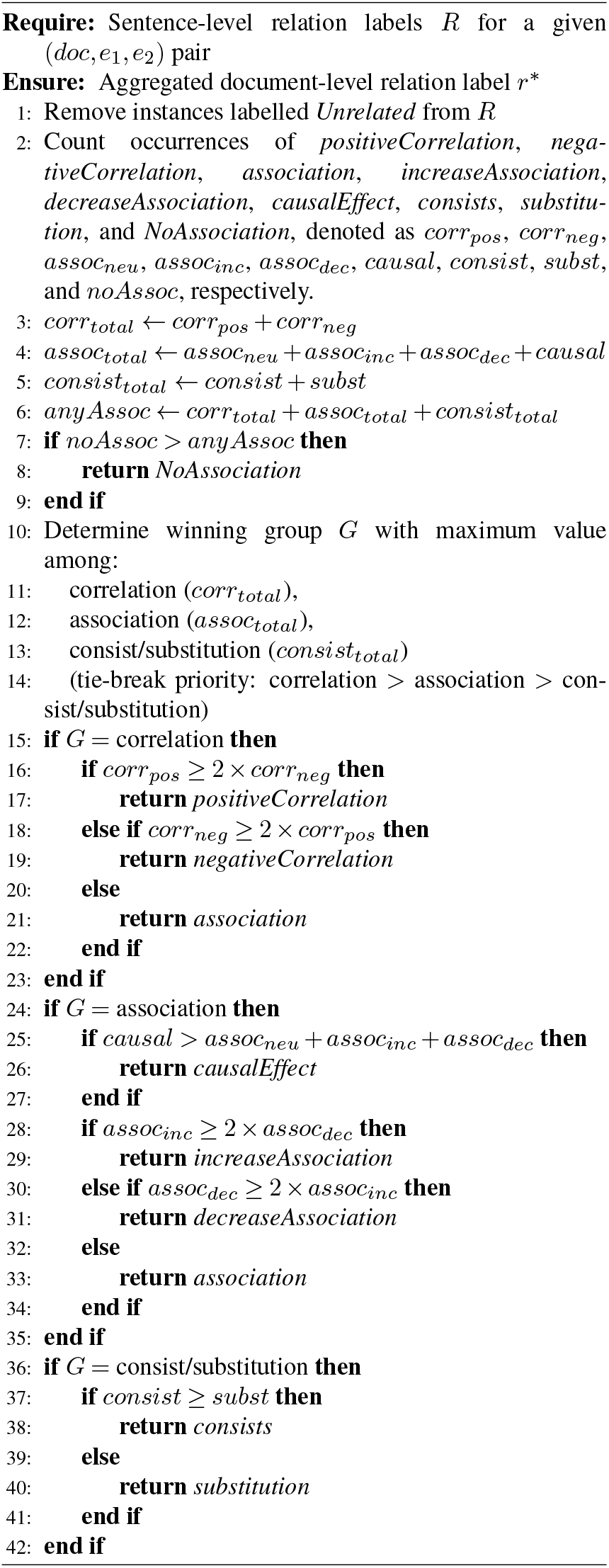

These document-level summarised extracted relations are then aggregated into two scores for each entity pair. The first is the Association Support (AS) score indicating the evidence for an association in general (Eqn. 1), and the second is an Effect Estimate (EE) score that indicates the direction (direct, inverse) of the association (Eqn. 2) using terminology from Figure 2 to indicate elements:

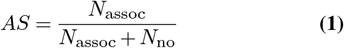

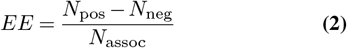

where *N*_pos_, *N*_neg_, and *N*_neu_ denote the number of documents whose aggregated relation corresponds to positive evidence (*increaseAssociation, positiveCorrelation*, or *consists*), negative evidence (*decreaseAssociation, negativeCorrelation*, or *substitution*), and neutral evidence (*causalEffect* or *association*), respectively. *N*_assoc_ = *N*_pos_ + *N*_neg_ + *N*_neu_, and *N*_no_ denotes the number of documents labelled as *NoAssociation*. All counts refer to document-level aggregated relations (with *Unrelated* excluded).

Eqn. 1(AS) signifies the grouping of direct, indeterminate, and inverse relations over total number of relations (including no association but excluding unrelated) to give a score limited between 0 and 1. Where Eqn. 2(EE) gives the direction of the combined evidence (direct associations minus inverse associations over the total number of associations) with the score ranging from -1 to 1. Here 0 indicates there is equal evidence for either side, and -1/+1 indicates all evidence is inverse/direct, respectively.

As an illustrative application of the CoCoS pipeline, we curated a subset of relations centered on key dietary factors. Specifically, we focus on relations between food-related entity mentions (e.g., `sugar-sweetened beverage` and `high-fat diet` (`HFD`), including `Western diet`) and disease/phenotype mentions (e.g., `insulin resistance/sensitivity/tolerance`, `type-2 diabetes`, `obesity/obese/overweight`, and `liver inflammation/injury`). Furthermore, we examine how both groups are associated with commonly reported proteins (`high-density lipoprotein` (`HDL`) `cholesterol,low-density lipoprotein` (`LDL`) `cholesterol`, and `alanine aminotransferase` (`ALT`)) and microbiota (*Firmicutes* and *Bacteroidetes*).

## Results

### Model selection (validation dataset)

We evaluated 14 LMs, each with 2-3 different strategies, using the training data, and selected the best-performing models for final test set evaluation. To ensure a fair and unbiased comparison, model selection was performed exclusively on the validation set based on the highest multi-class F1 score within each model architecture (BERT, LLM), training domain (general, biomedical), and training strategy (zero-shot, hierarchical, reflection). See the Supplementary Materials (Validation Results, Table S1) for the full results table. The top two models for the fine-tuned BERT architecture were *BiolinkBERT* (0.64) and *BioBERT* (0.64); for general-purpose LLMs, *gpt-oss-20B* (0.67) and *LLaMA3-70B* (0.64); and for biomedical domain-adapted LLMs, *Meerkat-70B* (0.66) and *MedGemma-27B* (0.64). The test set was reserved exclusively for final performance assessment to identify the most suitable modelling paradigm for diet–disease–bioentity relation extraction and was not used for model training, tuning, or selection.

### Multi-class classification (test dataset)

In our evaluation, *gpt-oss-20B* consistently outperformed all other models across accuracy, macro F1, precision, and recall on the test set, as shown in Table 2. It achieved an accuracy of 0.71, a macro F1 of 0.64, a precision of 0.69, and a recall of 0.63, marking the overall best performance among all compared models. The 20B-parameter model outperformed both biomedical-specific supervised models *BiolinkBERT* (accuracy 0.55, macro F1 0.52) and *BioBERT* (accuracy 0.49, macro F1 0.48), as well as larger domain-adapted generative models *MedGemma-27B* (accuracy 0.61, macro F1 0.61) and *Meerkat-70B* (accuracy 0.62, macro F1 0.57).

**Table 2.** Multi-class classification results for the test set. Top result for each metric is given in bold font, and second best is underlined. F1, precision and recall are reported as macro.

| Model name | Accuracy | F1 Score | Precision | Recall |
| --- | --- | --- | --- | --- |
| BioBERT | 0.4850 | 0.4768 | 0.5210 | 0.5131 |
| BiolinkBERT | 0.5492 | 0.5233 | 0.5353 | 0.5903 |
| Meerkat-70B | 0.6155 | 0.5712 | 0.5860 | <u>0.6018</u> |
| MedGemma-27B | 0.6126 | <u>0.6065</u> | <u>0.6706</u> | 0.5999 |
| LLaMA3-70B | <u>0.6287</u> | 0.5934 | 0.6293 | 0.5884 |
| gpt-oss-20B | **0.7106** | **0.6431** | **0.6917** | **0.6270** |

### Binary classification result (test dataset)

From an information extraction perspective, a key objective is to reliably distinguish whether a given entity pair should be extracted for downstream use or disregarded. To this end, we reformulated the task as a binary classification problem, grouping *Unrelated* and *NoAssociation* into the negative class and all remaining relation types into the positive class. In the binary classification setting, *gpt-oss-20B* achieved the strongest overall performance across all key metrics, as shown in Table 3. Specifically, its accuracy was 0.90, macro F1 0.92, precision 0.90, and recall 0.94, outperforming all other models by a clear margin. The next best model, *LLaMA3-70B*, obtained lower scores (accuracy 0.85, F1 0.89, precision 0.83) but having a larger recall of 0.97. The only category *gptoss-20B* did not achieve the best performance in was the binary recall for which *MedGemma-27B, BioBERT*, and *BiolinkBERT* all achieved 0.98.

**Table 3.** Binary classification results for the test set. Top result for each metric is given in bold font, and second best is underlined. F1, precision and recall are reported as macro.

| Model | Accuracy | F1 Score | Precision | Recall |
| --- | --- | --- | --- | --- |
| BioBERT | 0.7394 | 0.8312 | 0.7200 | <u>0.9830</u> |
| BiolinkBERT | 0.7674 | 0.8463 | 0.7438 | 0.9817 |
| Meerkat-70B | 0.8238 | 0.8715 | 0.8311 | 0.9161 |
| MedGemma-27B | 0.8353 | 0.8863 | 0.8065 | **0.9836** |
| LLaMA3-70B | <u>0.8501</u> | <u>0.8938</u> | <u>0.8313</u> | 0.9666 |
| gpt-oss-20B | **0.8967** | **0.9225** | **0.9038** | 0.9420 |

### Multi-document relation summarisation

Integrating the evidence across multiple documents and quantifying the associations between food-related items and health indicators (both disease phenotypes and bioentities) was done by calculating two scores: the Association Support (AS) score that reflects the overall evidence for any relationship, and the Effect Estimate (EE) score that indicates the direction of that relationship as either direct or inverse. This was visualised as a directed graph in Figure 3, with the corresponding Co-CoS scores for selected relations summarised in Table 4. The strongest evidence was found for `HFD` impacting on `LDL-cholesterol` with an EE score of 0.75, indicating that at least 75% of 28 documents (28/29 documents report any type of association, AS score = 0.97) indicate it is a direct relationship opposed to any other type. The relationship most prevalent in the corpus is `HFD` with `obesity/obese` with an EE of 0.71/0.65, with an AS of 0.96/1.00. The microbiota *Firmicutes* (direct) and *Bacteroidetes* (inverse) also have consistent EE scores with `HFD` (0.71/-0.73), `obese` (0.52/-0.58), and `obesity` (0.33/-0.50), but opposite with `overweight` (-0.13/0.29) with the low scores indicating the conflicting evidence for overweight opposed to obese groups.

**Fig. 3.**
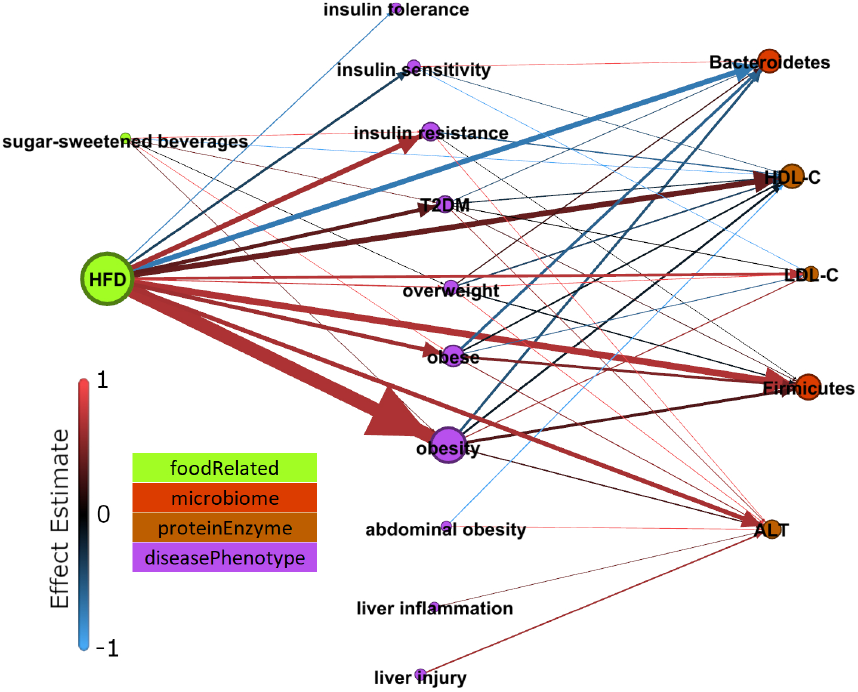
Graphical visualisation of CoCoS scores for a small set of relations across 4,445 documents. Node size is proportional to the number of documents each node is part of a relation. Node colour relates to the entity category. Edge width is proportional to the number of documents that reports the relation. Edge colour relates to the Effect Estimate (Eqn. 2).

**Table 4.**
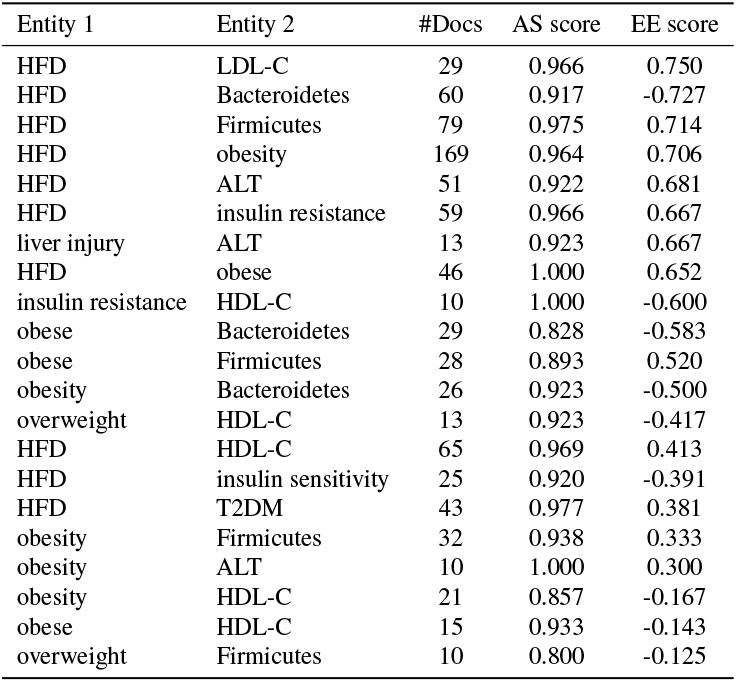
CoCos example (extracted from Figure 3) showing all scores for relations found in at least 10 documents. Association Support (AS) score (Eqn. 1) indicates the evidence found for an association (0 to 1), and the Effect Estimate (EE) score (Eqn. 2) indicates the directionality (-1 to 1). Abbreviations: HFD = high-fat diet, T2DM = type-2 diabetes mellitus, HDLC = high-density lipoprotein cholesterol, LDLC = low-density lipoprotein cholesterol, ALT = alanine aminotransferase.

## Discussion

We contributed a dedicated diet–disease-bioentity RE dataset covering 10 different relation types, including for the first time explicit negative examples. We evaluated both fine-tuned BERT- and LLM prompt-based models for their capabilities to capture complex relations between entities. We showed that generative LLMs, when guided with structured prompting strategies, hold strong potential for biomedical information extraction with an F1 over 0.90 for binary classification and multi-class classification of 0.64 F1.

Our evaluation reveals that generative models demonstrate significantly better generalisation compared to such supervised models. For instance, while the supervised *BioBERT* model dropped from a macro F1 of 0.64 on the validation set to 0.48 on the test set, our top model (*gpt-oss-20B*) maintained stable performance, achieving 0.67 and 0.64 on validation and test, respectively. This substantial gap indicates that prompt-based generative models exhibit far greater robustness to domain shifts and unseen examples than traditional supervised architectures, which tend to overfit both the training distribution and the hyperparameter optimisation process that is tightly tuned to the validation set. For example, we demonstrated that *DiMB-RE* was heavily dependent on the gold-standard labels within their benchmark, as it did not generalise to new data here. While our best model achieved an F1 of 0.64 (multi-class), this is in contrast to other existing methods that only scored 0.45 when evaluated under the same conditions (end-to-end) on their own corpus (12), but dropping to 0.29 when evaluated on our out-of-context data. Hence, these traditional supervised architectures remain sensitive to training distributions and validation-tuned hyperparameters, making prompting-based generative models more suitable as they achieve superior robustness and balanced precision–recall trade-offs.

In biomedical relation extraction, most models identify relationships between entities at the sentence or paragraph level. However, real-world evidence in biomedical literature is often distributed across multiple documents. To enable cross-document reasoning and synthesis we introduced CoCoS, a post-processing framework that aggregates relation extraction results for identical entity pairs observed across different contexts.

Unlike previous biomedical knowledge graph construction efforts such as PubMedKG (30), which primarily aggregate entities based on simple co-occurrence, our approach aggregates evidence at the document level to enable an automatic systematic review of diet–disease relationships along with bio-entities. Specifically, CoCoS is designed to summarise diverse and sometimes conflicting findings reported across multiple studies, where multiple independent relation extraction outputs are contextualised and merged into a coherent summary that reflects the overall scientific consensus or disagreement. CoCoS effectively summarised several relations that are well-known with the correct effect estimate (sign), such as obesity/obese and the high-fat diet with *Firmicutes* (direct) and *Bacteroidetes* (inverse) that reflect the association between obesity and the *Firmicutes*-to-*Bacteroidetes* ratio (31) and opposite signs of LDL-C (bad) and HDL-C (good) with different diseases and phenotypes consistent with the current body of knowledge (32). On the other hand, CoCoS did not find evidence beyond ‘*association*’ (1 document) between sugar-sweetened beverages and T2DM, while this is reported in the Burden of Proof (BoP) (10) resource as a weak direct effect.

However, the BoP only considers randomised controlled trials and relies on manual effort to extract data and perform the meta-analyses. Hence, only 88 of the 177 BoP entries relate to modifiable dietary or cardiometabolic risks, of which 48 relate to outcomes linked to metabolic syndrome. In contrast, our approach is fully computational and aims to bring together evidence across all study methodologies. In future work, we aim to release an extended version of CoCoS that integrates document-level weighting based on study methodology classification aligned to the hierarchy of evidence. This will bridge the gap between the fast computational approaches for systematic analysis and the time-intensive human systematic review process and generate testable hypotheses for future meta-analyses.

In summary, our results confirm that generative LLMs outperform supervised models for binary discrimination tasks, where the goal is to identify whether a relation exists between two entities, and extend well to multi-class RE. These models can be used to pre-select relevant bits of information for human review at a fraction of the time that it would take a human reader. Further improving these models will make their use alongside a human researcher in a systematic review a not-so-distant possibility.

## AUTHOR CONTRIBUTIONS

Conceptualization: MR, TB, and JP; Methodology: DC and JP; Software: DC, YG; Validation: DC, KZ, AL, DZ, and JP; Formal analysis: DC and YG; Investigation: DC and YG; Resources: MR, TB, and JP; Data Curation: DC, KZ, AL, DZ, and JP; Writing - Original Draft: DC and JP; Writing - Review & Editing: DC, YG, AL, TB, and JP; Visualization: DC; Supervision: JP; Project administration: JP; Funding acquisition: MR, TB and JP. All authors that contributed beyond data curation have read and agreed to the published version of the manuscript.

For more information see https://www.codiet.eu/.

## Supplementary Note 1: Models

We evaluate two categories of pretrained language models: **general-purpose LMs**, which serve as baselines across diverse NLP tasks, and **biomedical LMs**, which are specifically adapted to biomedical and clinical corpora. For further comparison, we apply the relation extraction pipeline from DiMB-RE on our diet-disease dataset to evaluate model performance, using a method originally developed for diet-microbiome relations.

In information extraction (IE) tasks, traditional BERT-like models trained with supervised learning remain the dominant paradigm, as they are effective for structured relation extraction and entity-level classification. However, due to their potential for broader generalisation, generative large language models have recently gained increasing attention. Accordingly, we evaluated both well-established BERT-based models and state-of-the-art generative models that could be locally deployed within our available GPU specifications.

For model deployment, we used HuggingFace, vLLM, and LLaMA.cpp ^1^ implementations, and applied quantisation and optimisation techniques tailored to the NVIDIA L40S GPU. Below, we summarise the models considered in each category. For the General-purpose LMs, we utilise the below models.

1. **BERT-base**: A standard Transformer-based model widely regarded as a strong baseline in general NLP.
2. **LLaMA-8B**: A medium-scale (8B parameters) model from the LLaMA family, capable of general text understanding and generation.
3. **LLaMA-70B**: A large-scale model (70B parameters) with strong few-shot reasoning ability.
4. **Qwen-32B**: A multilingual model developed by Alibaba, achieving competitive results on general and reasoning tasks.
5. **DeepSeek-Qwen-32B**: A distilled variant of DeepSeek-R1 built upon Qwen-32B, designed for improved efficiency–performance trade-offs.
6. **DeepSeek-LLaMA-70B (q4_k_s)**: A quantised and distilled variant of DeepSeek-R1 built upon LLaMA-70B, enabling memory-efficient inference while maintaining strong reasoning performance. We use llama.cpp to utilise this model.
7. **gpt-oss:20B**: An open-weight reasoning model based on a mixture-of-experts Transformer architecture, trained through large-scale distillation and reinforcement learning, achieving strong general-purpose performance with notable strengths in reasoning and related tasks. We use vllm to utilise this model.

Also, we utilised Biomedical LMs. We included a range of biomedical and clinical domain–adapted models to assess their performance in comparison with recent generative large language models that have been specialised for biomedical or medical text understanding.

- **BioBERT**: A domain-adapted BERT model pretrained on large-scale biomedical corpora, including PubMed abstracts and PMC articles.
- **PubMedBERT**: A biomedical-specific BERT trained exclusively on PubMed abstracts, yielding strong domain performance.
- **BiolinkBERT**: A biomedical BERT variant aligned with knowledge graphs, effective for entity linking and relation extraction.
- **Meerkat-8B**: A lightweight biomedical LLM (8B parameters) specialised in clinical and biomedical text processing.
- **Meerkat-70B**: A large-scale biomedical LLM (70B parameters), designed for complex reasoning in biomedical and clinical domains.
- **MedGemma-27B**: An instruction-tuned biomedical model (27B parameters) from Google’s Gemma family, tailored for clinical QA and instruction following.

## Supplementary Note 2: Data splitting and evaluation metrics

For the multi-class classification setting, we report standard metrics including accuracy, and macro-averaged precision, recall, and F1-score. Macro averaging is used to account for label imbalance across classes. In addition, we also conduct a binary classification evaluation. Since one of our main goals is to construct a knowledge graph, it is essential to determine whether a given entity pair is related in any way. For this purpose, we group *NoAssociation* and *Unrelated* into the negative class, and all other relation types into the positive class. For this binary task, we again evaluate using accuracy and macro-averaged precision, recall, and F1-score.

## Supplementary Note 3: Validation Results

### Multi-class classification on the validation set

The validation set results in Table S1 show that the performance varies considerably across architectures and inference strategies. We therefore selected the top two models in terms of macro F1 from each category (general LMs, biomedical LMs, and generative LLMs). These selected models were then carried forward to the independent test set evaluation. This design ensured a fair comparison between traditional supervised models and large generative LLMs under different prompting strategies, while preventing overfitting to a single inference style.

Our proposed *Hierarchical prompt + reflection* strategy applied to *gpt-oss-20B* achieved the highest macro F1-score (0.6731), confirming the effectiveness of this approach in handling complex multi-class relation extraction. This result highlights the advantage of combining structured decision prompting with reflective reasoning for challenging biomedical relations, even when using a locally deployable generative LLM.

**Fig. S1.**
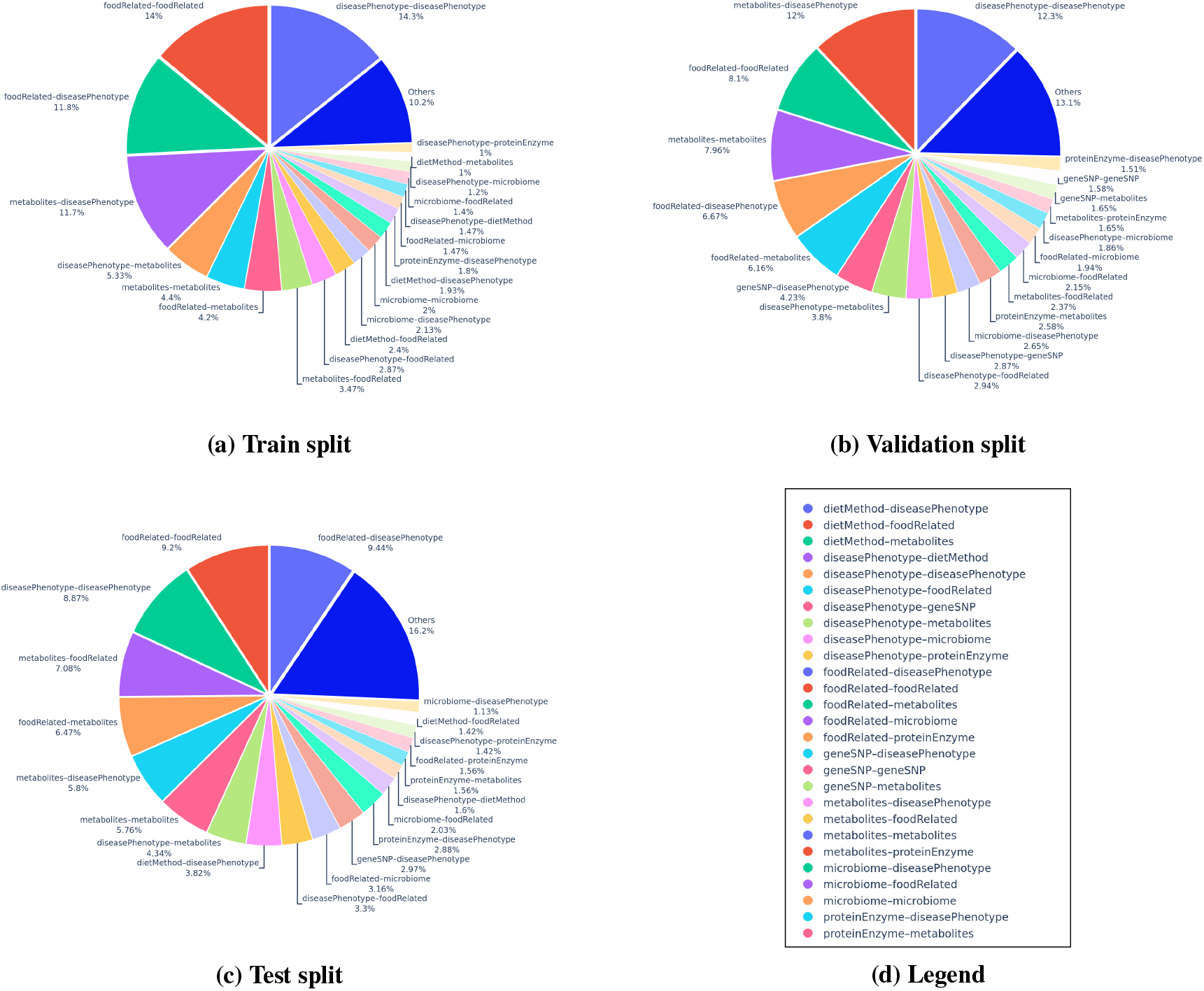
Top 20 entity pair distributions across splits (train/val/test). Each pie chart shows the percentage composition of the most frequent top pairs. The bottom-right panel is a placeholder for legend or extended example.

Our proposed *gpt-oss-20B* model with the *Hierarchical prompt + reflection* strategy demonstrated strong generalisation capability, achieving the highest macro F1-score (0.6731). This result indicates that prompt-based generative LLMs can capture underrepresented relation types more effectively, thereby overcoming the data distribution bias often observed in supervised learning. Nevertheless, the BERT-based supervised models also showed notable strengths. Despite being considerably smaller and computationally lighter, they achieved competitive accuracy and precision compared to large generative models. For instance, the *DiMB-RE* model obtained the highest overall accuracy (0.7238), outperforming our best-performing generative model in that metric (*gpt-oss-20B*, 0.6741).

The proposed *Hierarchical prompting* strategy proved particularly effective for generative LLMs, leading to consistent performance improvements across most models. In contrast, it resulted in performance degradation in all BERT-based models, suggesting that hierarchical decision-making is not a universal performance booster. Instead, it is especially beneficial for generative models when properly designed to mitigate context-length limitations and enhance reasoning depth through structured decomposition.

Domain adaptation benefits traditional supervised BERT models but not modern generative LLMs. In the BERT-based group, domain-adapted biomedical LMs like BioBERT, PubMedBERT outperformed general models, indicating that domain adaptation remains advantageous for supervised learning with BERT-like LMs. In contrast, the general-purpose *gpt-oss-20B* significantly outperformed domain-specific biomedical LLMs such as *Meerkat-70B* and *MedGemma-27B*. This suggests that modern instruction-tuned generative models are less dependent on domain-specific corpora and rely more on their reasoning and instruction-following capabilities. Biomedical generative LLMs are often validated for question answering rather than information extraction, which may explain their relatively weaker performance in this task.

**Table S1.** Multi-class classification results for the validation set. The validation set was used to select the best models and approaches to evaluate with the independent test set based on the F1-score. The top model from each category (BERT-based, general LLM, biomedical LLM) for each metric is indicated in bold font, with the second best models underlined. We omit DiMB-RE with the hierarchical experts approach due to difficulties in modifying the original codebase.

| BERT model name | Type | Accuracy | F1 (Macro) | Precision (Macro) | Recall (Macro) |
| --- | --- | --- | --- | --- | --- |
| Bert-base | Normal | 0.6307 | 0.5224 | 0.5466 | 0.5210 |
| BioBERT | Normal | 0.7026 | <u>0.6358</u> | <u>0.6851</u> | **0.6171** |
| PubmedBERT | Normal | 0.7083 | 0.6276 | 0.6667 | 0.6145 |
| BiolinkBERT | Normal | <u>0.7147</u> | **0.6440** | **0.8006** | <u>0.6149</u> |
| DiMB-RE | Normal | **0.7238** | 0.2940 | 0.4262 | 0.2360 |
| Bert-base | Hierarchical - Individual Experts | 0.5326 | 0.3980 | 0.4500 | 0.4084 |
| BioBERT | Hierarchical - Individual Experts | 0.6158 | 0.4686 | 0.4813 | 0.4640 |
| PubmedBERT | Hierarchical - Individual Experts | 0.6724 | 0.5017 | 0.5085 | 0.5010 |
| BiolinkBERT | Hierarchical - Individual Experts | 0.6330 | 0.4613 | 0.4954 | 0.4655 |
| DiMB-RE | Hierarchical - Individual Experts | - | - | - | - |
| General LLM name | Type | Accuracy | F1 (Macro) | Precision (Macro) | Recall (Macro) |
| Llama-3.1-8B | Zero-shot | 0.3980 | 0.4268 | **0.7405** | 0.3922 |
| Qwen2.5-32B | Zero-shot | 0.5887 | 0.5927 | 0.6127 | <u>0.6584</u> |
| DeepSeek-R1-Distill-Qwen-32B | Zero-shot | 0.6365 | 0.5983 | 0.6316 | 0.6510 |
| DeepSeek-R1-Distill-Llama-70B | Zero-shot | 0.5920 | 0.5990 | 0.6632 | 0.6285 |
| Llama-3.3-70B-Instruct | Zero-shot | 0.5712 | 0.5447 | 0.5781 | 0.5962 |
| gpt-oss-20b | Zero-shot | 0.6303 | 0.6454 | 0.6829 | 0.6674 |
| Llama-3.1-8B | Hierarchical prompt | 0.5626 | 0.4618 | 0.4997 | 0.4680 |
| Qwen2.5-32B | Hierarchical prompt | 0.6065 | 0.5883 | 0.6979 | 0.5957 |
| DeepSeek-R1-Distill-Qwen-32B | Hierarchical prompt | 0.6052 | 0.5834 | 0.6921 | 0.5896 |
| DeepSeek-R1-Distill-Llama-70B | Hierarchical prompt | <u>0.6233</u> | 0.5811 | 0.6621 | 0.5997 |
| Llama-3.3-70B-Instruct | Hierarchical prompt | 0.5763 | 0.6018 | 0.6599 | 0.6264 |
| gpt-oss-20b | Hierarchical prompt | 0.6210 | 0.6380 | 0.6557 | 0.6749 |
| Llama-3.1-8B | Hierarchical prompt + reflection | 0.5793 | 0.4832 | 0.5378 | 0.4821 |
| Qwen2.5-32B | Hierarchical prompt + reflection | 0.6137 | 0.5910 | <u>0.7156</u> | 0.5914 |
| DeepSeek-R1-Distill-Qwen-32B | Hierarchical prompt + reflection | <u>0.6413</u> | 0.5954 | 0.6309 | 0.6451 |
| DeepSeek-R1-Distill-Llama-70B | Hierarchical prompt + reflection | 0.6370 | 0.5871 | 0.6840 | 0.5913 |
| Llama-3.3-70B-Instruct | Hierarchical prompt + reflection | 0.6259 | <u>0.6423</u> | 0.7118 | 0.6377 |
| gpt-oss-20b | Hierarchical prompt + reflection | **0.6741** | **0.6731** | 0.6885 | **0.6939** |
| Biomedical LLM name | Type | Accuracy | F1 (Macro) | Precision (Macro) | Recall (Macro) |
| meerkat-8b-v1.0 | Zero-shot | 0.3877 | 0.3546 | 0.6150 | 0.3812 |
| meerkat-70b-v1.0 | Zero-shot | 0.5671 | 0.6186 | <u>0.6907</u> | 0.6266 |
| medgemma-27b-it | Zero-shot | 0.6099 | <u>0.6438</u> | **0.7530** | <u>0.6213</u> |
| meerkat-8b-v1.0 | Hierarchical prompt | 0.4741 | 0.4069 | 0.4984 | 0.4185 |
| meerkat-70b-v1.0 | Hierarchical prompt | 0.5871 | 0.6129 | 0.6331 | 0.6598 |
| medgemma-27b-it | Hierarchical prompt | 0.6640 | 0.5792 | 0.6060 | 0.5785 |
| meerkat-8b-v1.0 | Hierarchical prompt + reflection | 0.4579 | 0.3578 | 0.5005 | 0.3668 |
| meerkat-70b-v1.0 | Hierarchical prompt + reflection | <u>0.6468</u> | **0.6600** | 0.6784 | **0.6897** |
| medgemma-27b-it | Hierarchical prompt + reflection | **0.6698** | 0.6263 | 0.7230 | 0.6014 |

Finally, we observe consistent gains when applying our proposed reflection mechanism in generative LLMs. For the *gpt-oss-20B* model, both hierarchical prompting and the reflection step contributed to incremental improvements in F1 and recall, confirming that structured multi-stage reasoning and self-reflection enable more stable and accurate decision-making in complex relation extraction tasks.

## Supplementary Note 4: Prompts

This section provides the full prompts used at each stage of the RECoDe relation extraction decision flow (Fig. 2). Each prompt corresponds to a specific classifier (C1–C7) in the staged framework and implements the branching logic defined in the decision process.

**Fig. S2.**
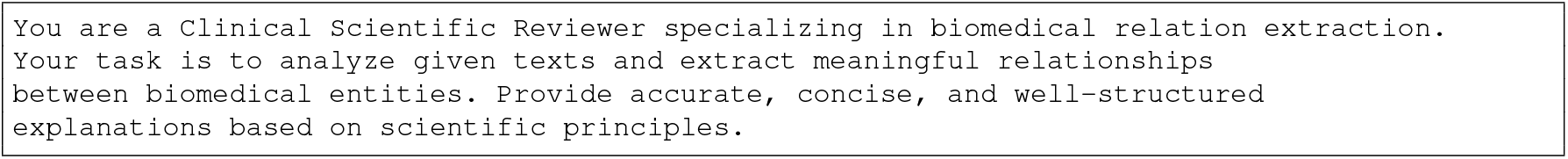
Global system instruction used to initialise the RECoDe relation extraction decision flow (Fig. 2). This prompt establishes the clinical reviewer role and defines the scientific reasoning context that guides all subsequent staged classification prompts (C1–C7) within the hierarchical RECoDe decision framework.

**Fig. S3.**
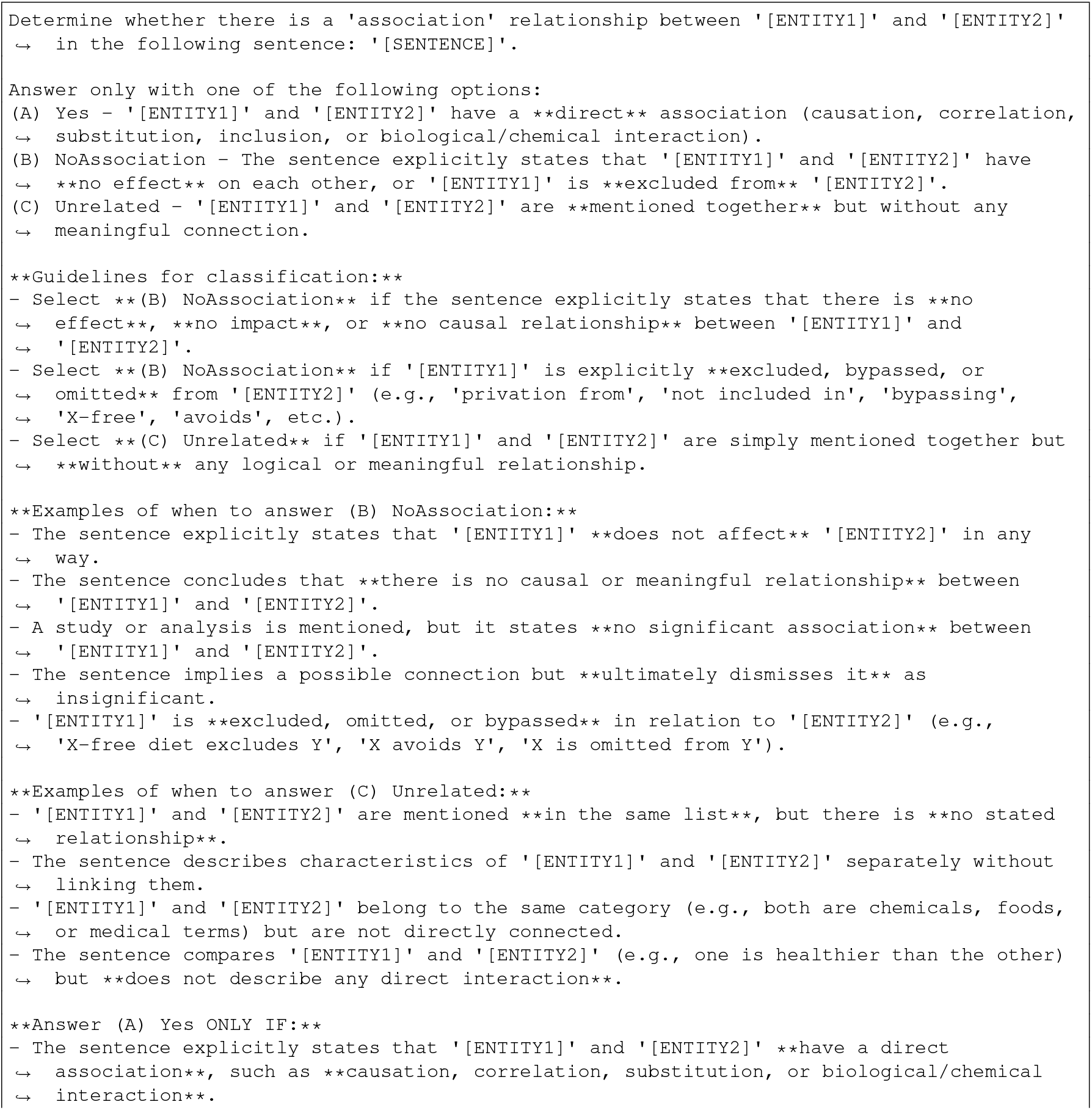
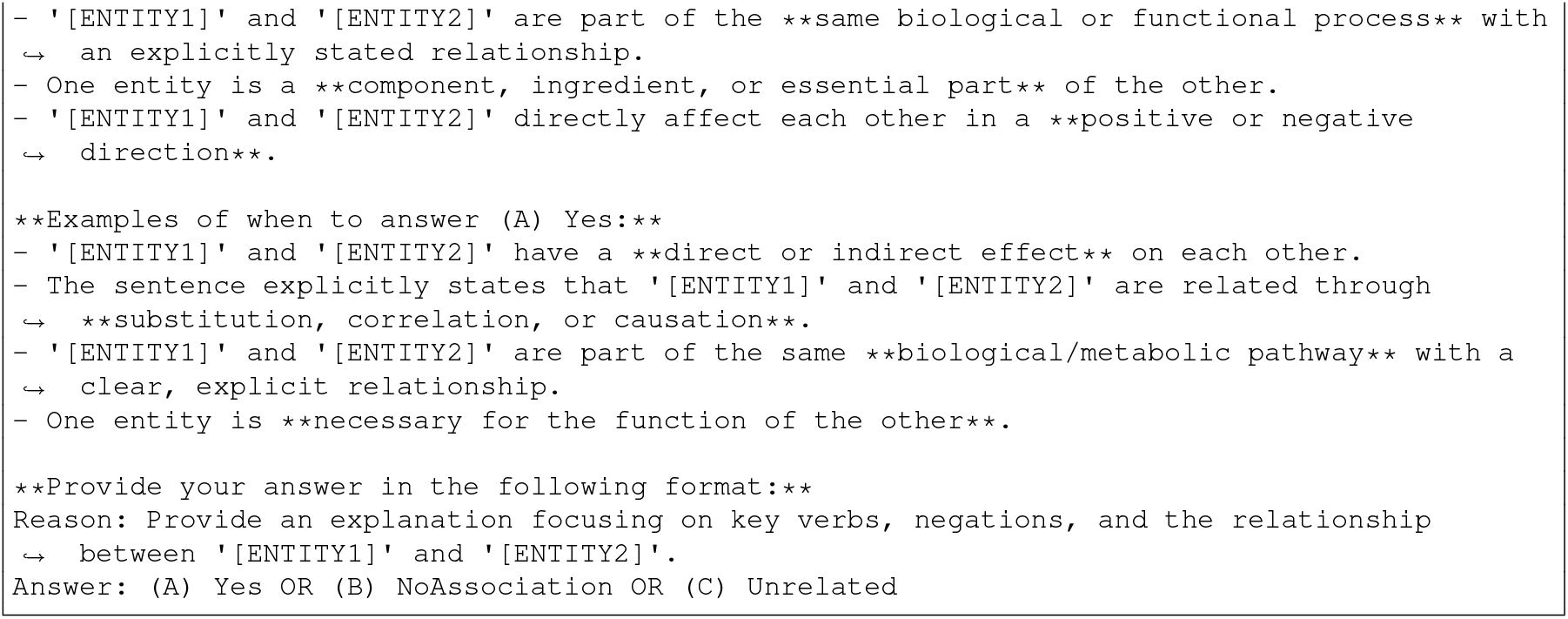
Prompt used in the C1First stage of the RECoDe decision flow (Fig. 2) to separate negative cases (NoAssociation and Unrelated) from positive relations, where Yes predictions are mapped to the C2Asso branch.

**Fig. S4.**
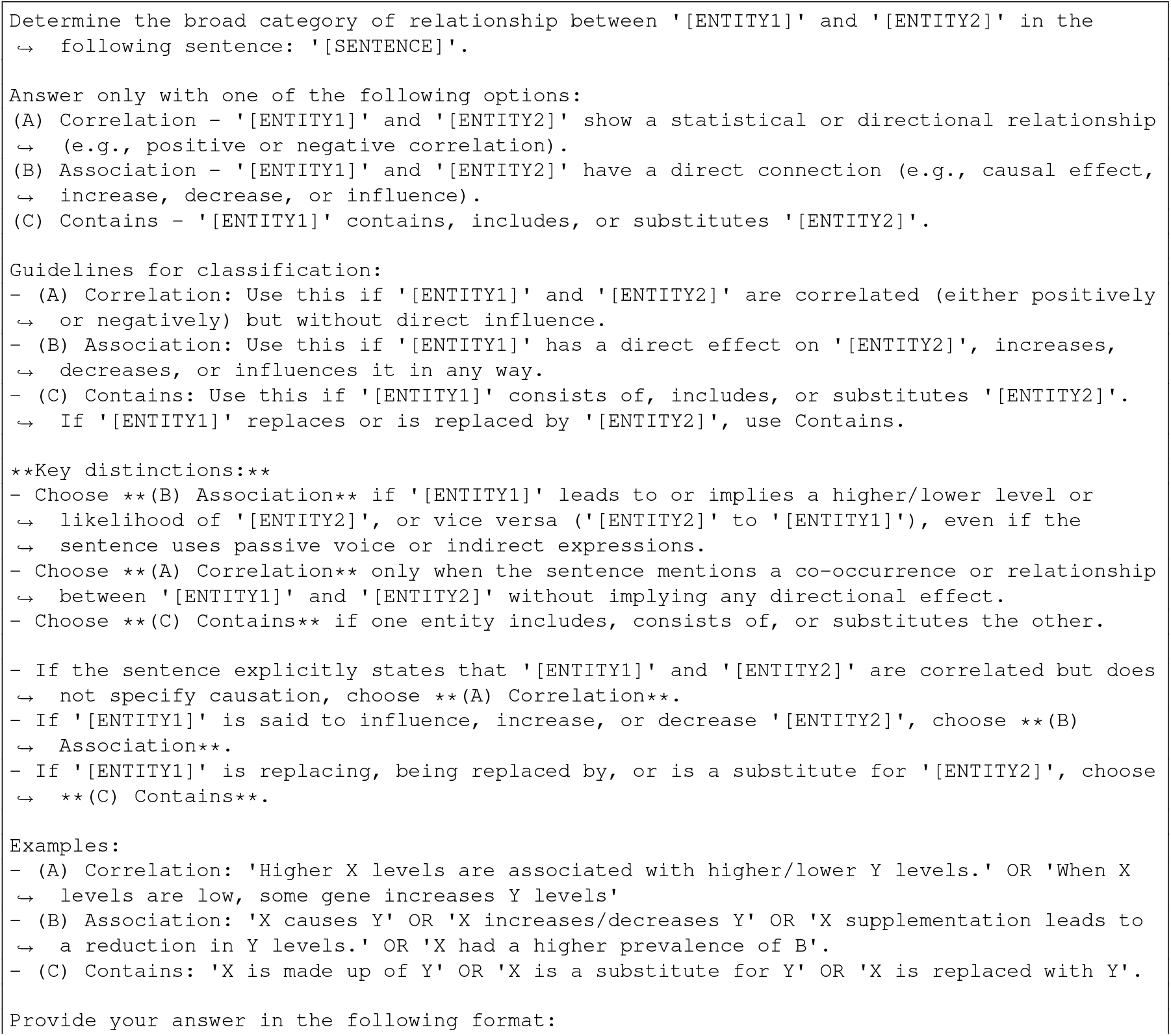
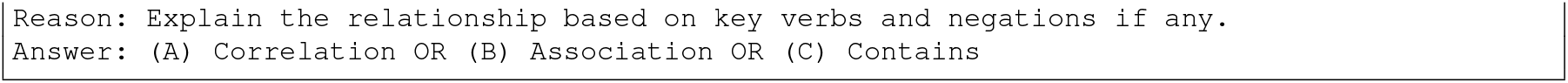
Prompt used in the C2Asso stage of the RECoDe decision flow (Fig. 2) to categorise relations from C1First into Correlation, Association, or Contains branches, which are subsequently routed to the C3Corr, C4MiddleAsso, and C6ContainSubsti stages, respectively.

**Fig. S5.**
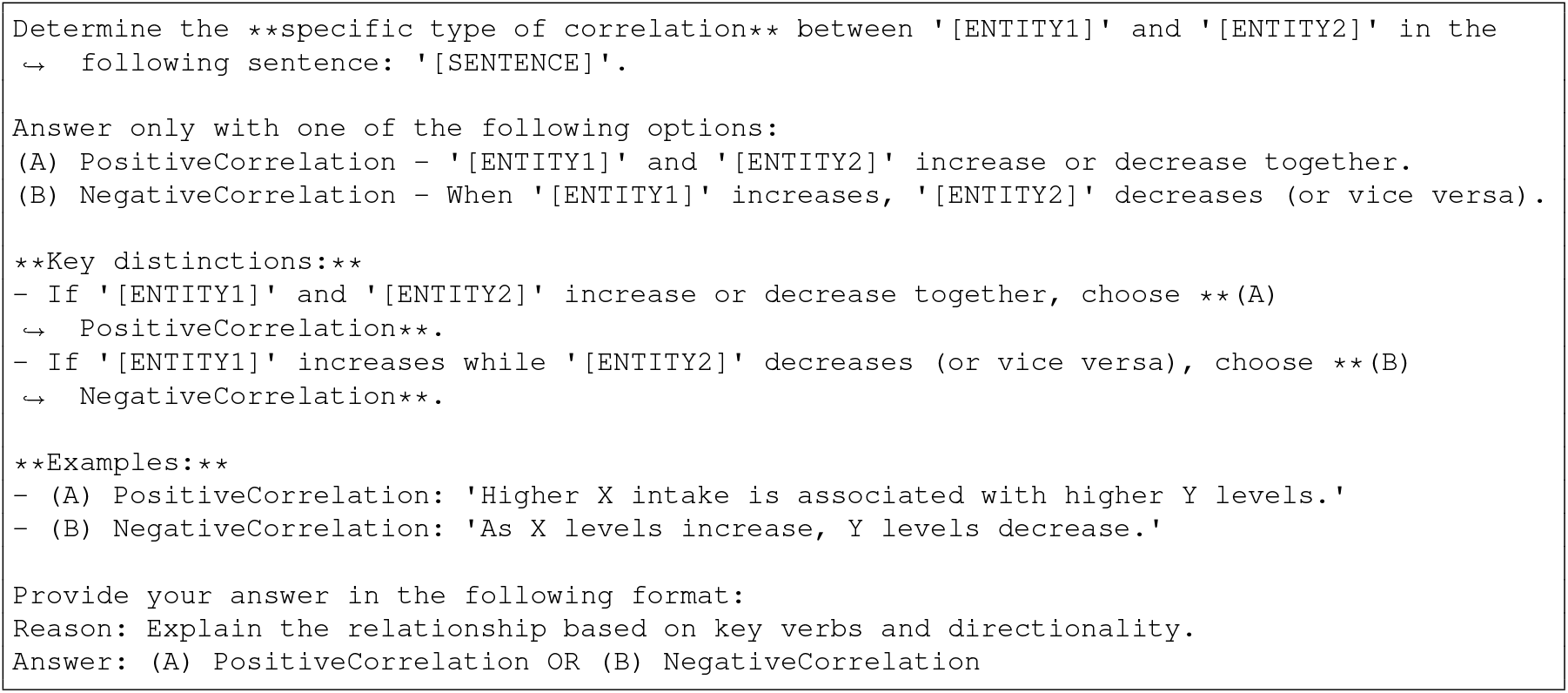
Prompt used in the C3Corr stage of the RECoDe decision flow (Fig. 2) to distinguish between PositiveCorrelation and NegativeCorrelation, with PositiveCorrelation predictions routed to the C7PositiveCorrReflection stage. The decision flow uses lowercase labels for consistency with the annotation schema, whereas the prompt presents capitalised forms as model output options.

**Fig. S6.**
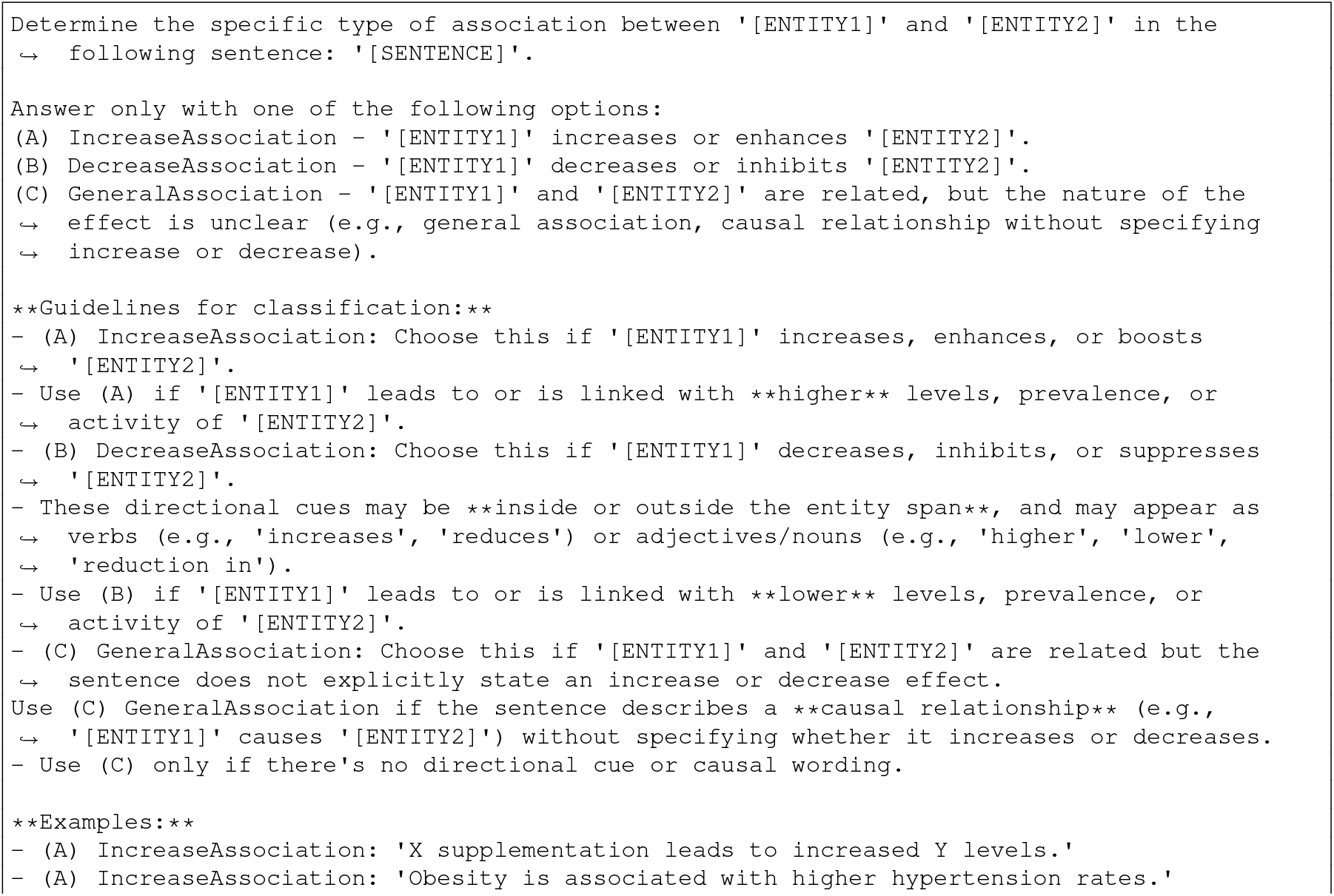
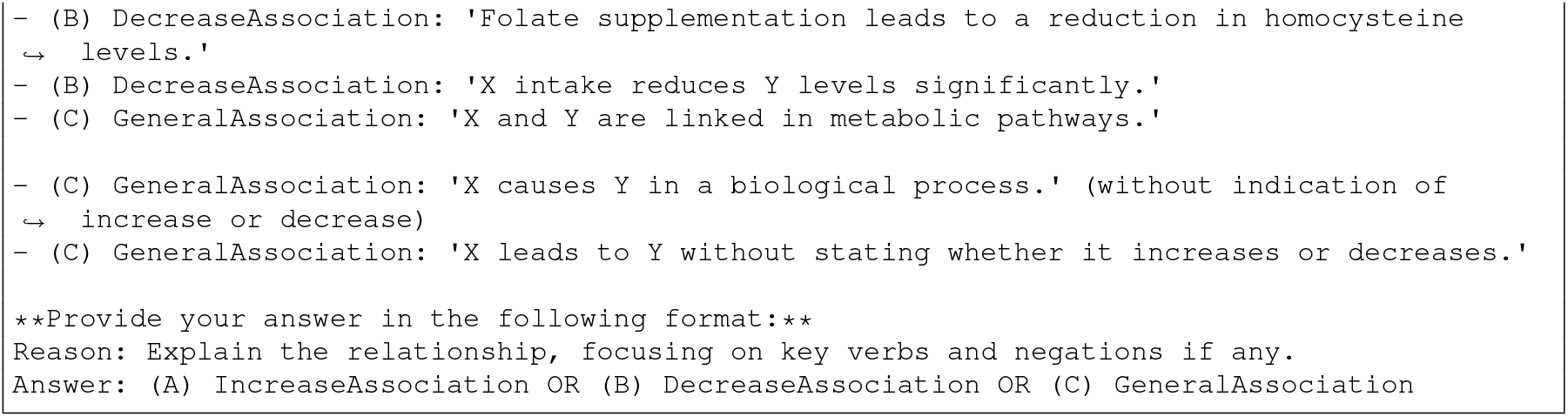
Prompt used in the C4MiddleAsso stage of the RECoDe decision flow (Fig. 2) to classify association relations into IncreaseAssociation, DecreaseAssociation, or GeneralAssociation, where GeneralAssociation predictions are routed to the C5AssoCausal stage. The decision flow uses lowercase labels for consistency with the annotation schema, whereas the prompt presents capitalised forms as model output options.

**Fig. S7.**
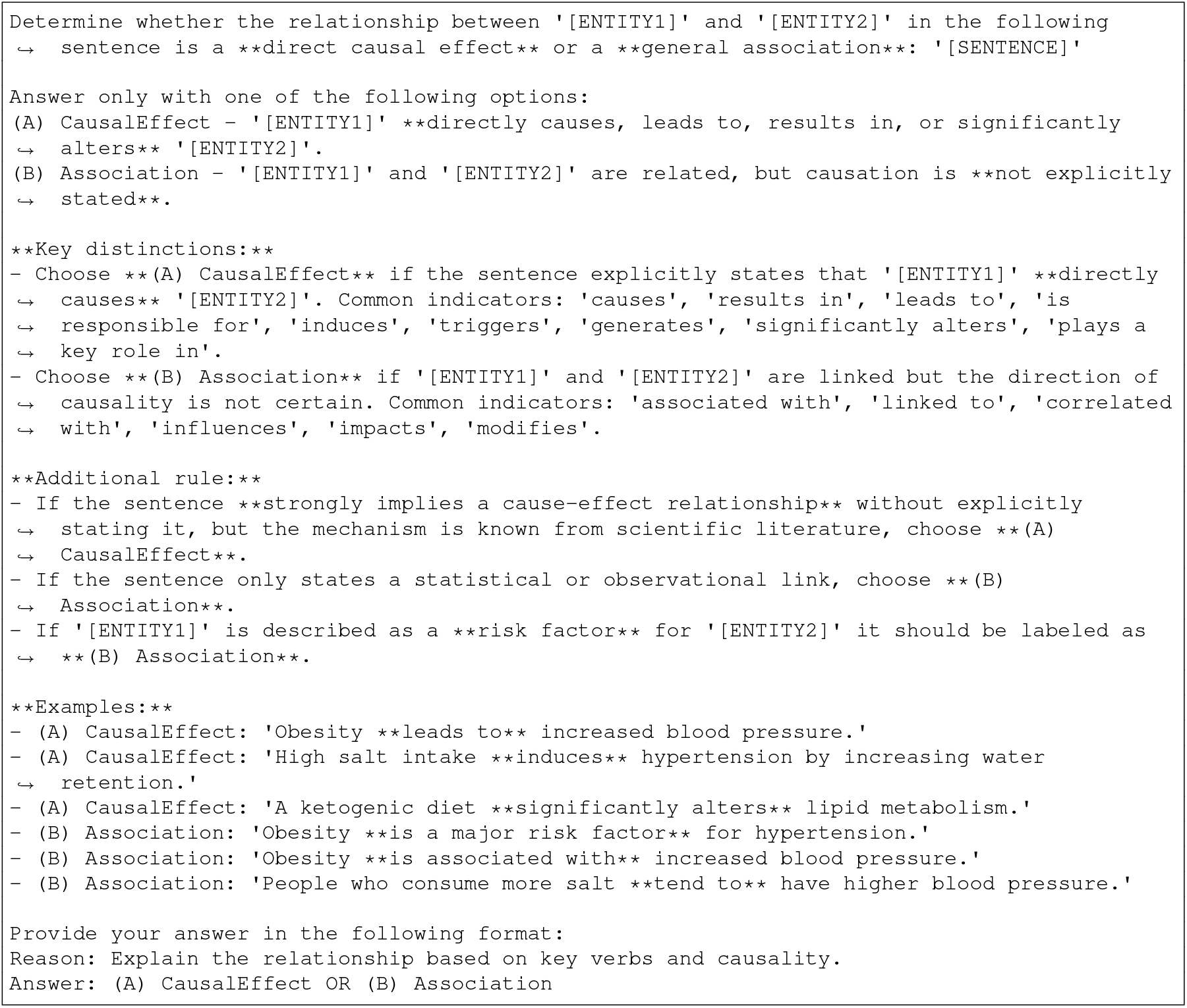
Prompt used in the C5AssoCausal stage of the RECoDe decision flow (Fig. 2) to distinguish between CausalEffect and Association for relations routed from the C4MiddleAsso stage. The decision flow uses lowercase labels for consistency with the annotation schema, whereas the prompt presents capitalised forms as model output options.

**Fig. S8.**
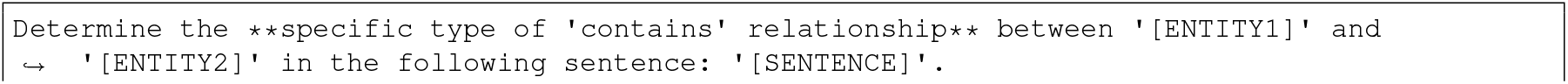
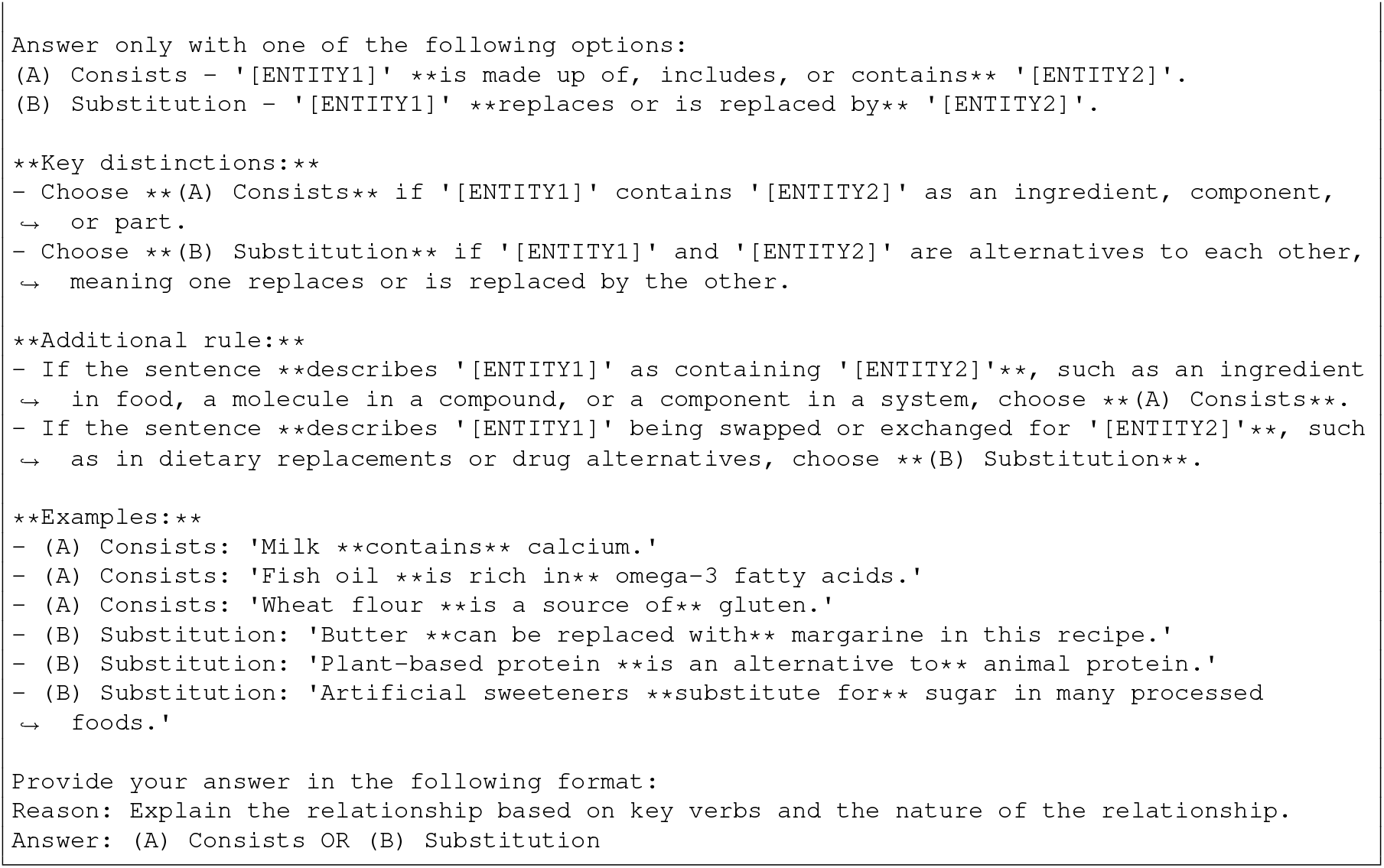
Prompt used in the C6ContainSubsti stage of the RECoDe decision flow (Fig. 2) to distinguish between Consists and Substitution relations for cases routed from the C2Asso Contains branch. The decision flow uses lowercase labels for consistency with the annotation schema, whereas the prompt presents capitalised forms as model output options.

**Fig. S9.**
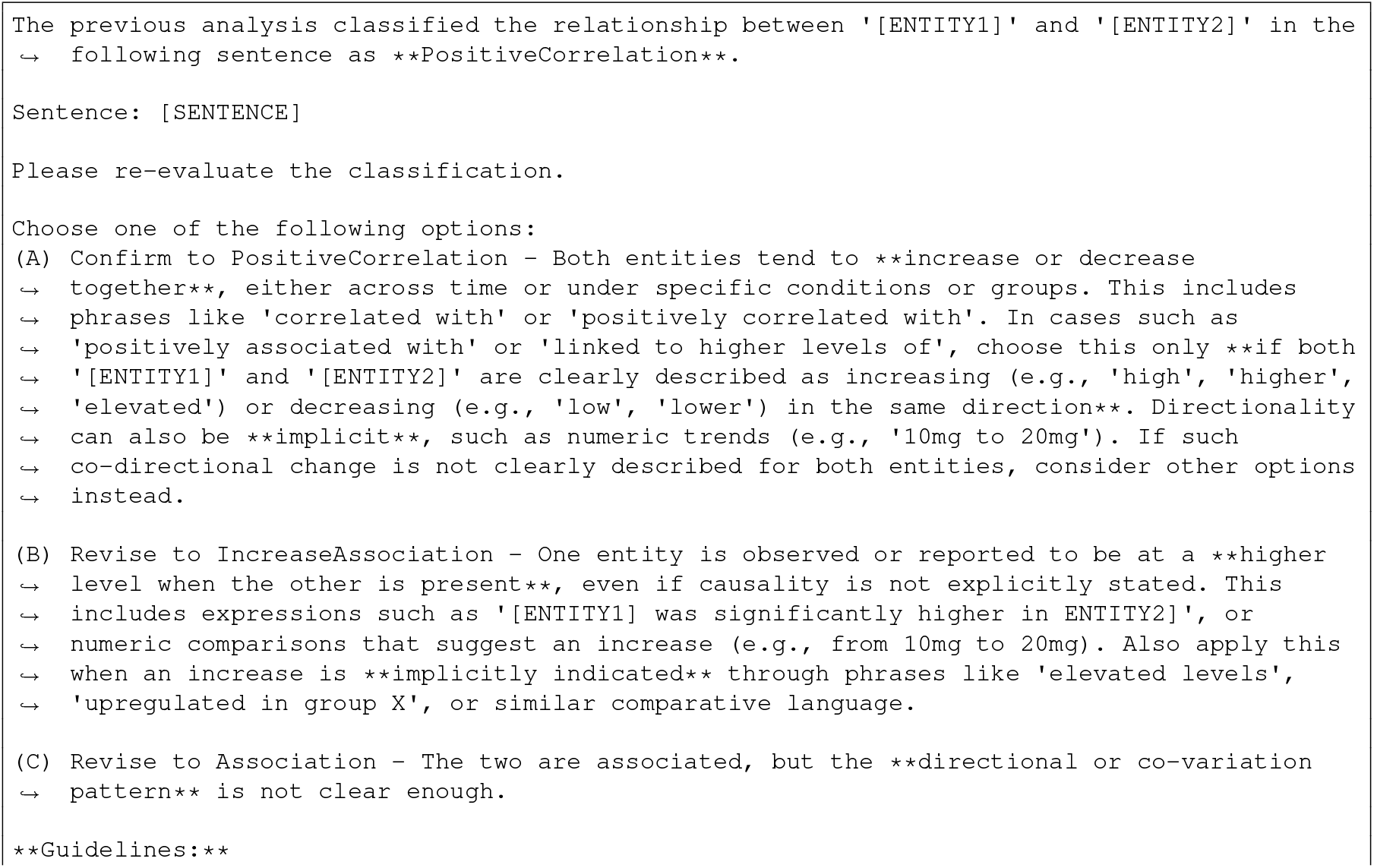
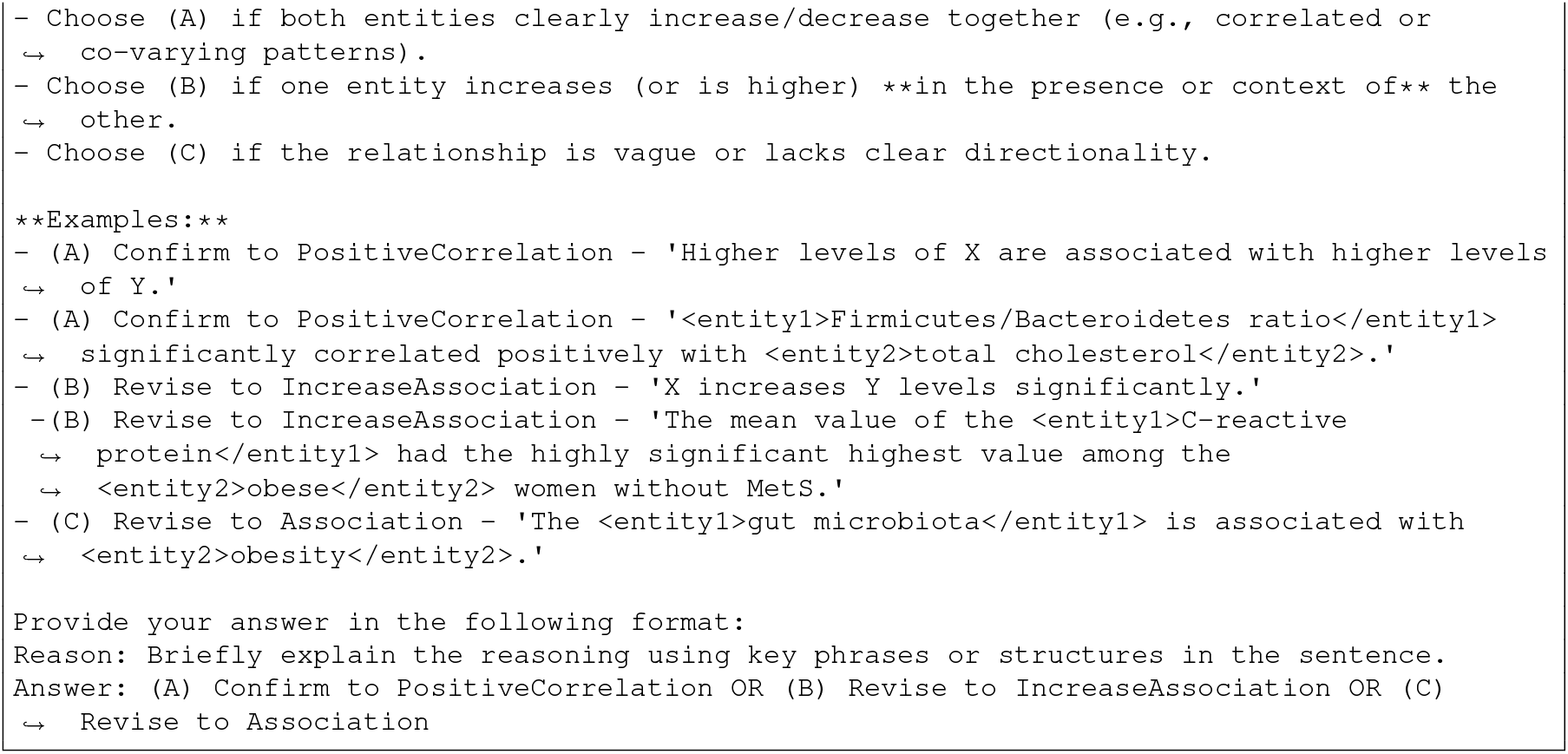
Reflection prompt used in the C7PositiveCorr stage of the RECoDe decision flow (Fig. 2) to re-evaluate PositiveCorrelation predictions, confirming PositiveCorrelation or revising them to IncreaseAssociation or Association when appropriate. The decision flow uses lowercase labels for consistency with the annotation schema, whereas the prompt presents capitalised forms as model output options.

## Footnotes

1 https://github.com/ggml-org/llama.cpp

